# EMPIAR: The Electron Microscopy Public Image Archive

**DOI:** 10.1101/2022.10.04.510785

**Authors:** Andrii Iudin, Paul K. Korir, Sriram Somasundharam, Simone Weyand, Cesare Cattavitello, Neli Fonseca, Osman Salih, Gerard J. Kleywegt, Ardan Patwardhan

**Affiliations:** European Molecular Biology Laboratory, European Bioinformatics Institute (EMBL-EBI), Wellcome Genome Campus, Hinxton, Cambridgeshire, CB10 1SD, United Kingdom

## Abstract

Public archiving in structural biology is well established with the Protein Data Bank (PDB; wwPDB.org) catering for atomic models and the Electron Microscopy Data Bank (EMDB; emdb-empiar.org) for 3D reconstructions from cryo-EM experiments. Even before the recent rapid growth in cryo-EM, there was an expressed community need for a public archive of image data from cryo-EM experiments for validation, software development, testing and training. Concomitantly, the proliferation of 3D imaging techniques for cells, tissues and organisms using volume EM (vEM) and X-ray tomography (XT) led to calls from these communities to publicly archive such data as well. EMPIAR (empiar.org) was developed as a public archive for raw cryo-EM image data and for 3D reconstructions from vEM and XT experiments and now comprises over a thousand entries totalling over 2 petabytes of data. EMPIAR resources include a deposition system, entry pages, facilities to search, visualise and download datasets, and a REST API for programmatic access to entry metadata. The success of EMPIAR also poses significant challenges for the future in dealing with the very fast growth in the volume of data and in enhancing its reusability.

## Introduction

In the past two decades we have witnessed rapid growth in the development of 3D electron microscopy (3DEM) and its use in biology. In structural biology, cryo-EM has supplanted X-ray crystallography as the method of choice for studying macromolecular structure (1–3) and techniques such as electron cryo-tomography and volume EM (vEM) enable the study of cells, tissues and in some cases even whole organisms with unprecedented levels of 3D ultrastructural detail (4–7).

A common theme across the 3DEM landscape is that the inherently noisy nature of biological EM images necessitates the collection of large amounts of image data and sophisticated image processing in order to obtain 3D reconstructions that can be interpreted in terms of biology. It is not uncommon for cryo-EM practitioners, when given the same raw dataset, to obtain different 3D reconstructions despite using the same (or similar) software (8,9) due to different choices of start-up models, parameters and workflows. Differences in specimen-preparation techniques and imaging may further exacerbate the problem. While it is possible to prepare samples and obtain a 3D reconstruction of a well-established and well-behaved sample in under a day (10), most experiments will take months if not years to complete and will involve significant resources in terms of high-end microscopy instrumentation and computing.

The potential variability of results and the cost in terms of time and resources present a strong case for the public archiving of 3DEM data as it enables reuse and validation. The Protein Data Bank (PDB) archive for atomic coordinate models of biological molecules was founded in 1971, making it the oldest surviving molecular archive in biology (11). In fact, the PDB and the institutes operating it (nowadays united in the Worldwide Protein Data Bank, wwPDB, wwPDB.org, (11–13)) have done much to cultivate a strong awareness of the need and benefits of, and community-wide support for public archiving in structural biology. They have served as a focal point, catalyst and facilitator on several issues requiring community consensus and support such as formats, data models, data standards, and validation. They have also inspired other communities to address public archiving of data. The Electron Microscopy Data Bank (EMDB; https://emdb-empiar.org) was founded in 2002 by the European Bioinformatics Institute (EMBL-EBI) with support from the cryo-EM community as an archive for 3D reconstructions derived from cryo-EM experiments (14). Nowadays, EMDB is both a partner in and a core archive of the wwPDB organisation and is growing exponentially, having doubled in size from 10,000 entries in February 2020 to 20,000 in May 2022.

While the 3D reconstructions archived in EMDB represent substantial value in terms of reusability, being the result of a complex process, they are of limited use in terms of validation. Public access to the original image data used to obtain a given reconstruction archived in EMDB would substantially facilitate validation. A key recommendation from both the 2010 Electron Microscopy Validation Task Force Meeting (15) and the 2011 “Data Management Challenges in 3D Electron Microscopy” (DMCEM, (16)) expert workshop was to establish a pilot image archive as a source of test data for methods development and to support validation of deposited reconstructions. Similar requests were made at community meetings such as the 3DEM Gordon Research Conferences (3DEM-GRC) and other expert workshops (17,18). Development of a proof-of-principle resource, which would evolve into EMPIAR (https://empiar.org/, (19)), began in 2012, leveraging EMBL-EBI’s IT infrastructure and expertise in archiving, distributing and providing high-speed access to big sequence datasets. EMPIAR’s inception coincided in time with the introduction of direct electron detectors in cryo-EM and the archive was in operation when the ensuing “resolution revolution” began to impact the field (20). This fortuitous timing meant that EMPIAR was able to play its part in and helped catalyse that revolution. The first entry (EMPIAR-10002; deposited by Sjors Scheres and released in 2013) was of the *S. cerevisiae* 80S ribosome, a dataset underpinning what was then the highest resolution ribosome structure in EMDB (~4Å) and demonstrating the merits of motion correction and frame averaging of direct electron detector images (21). Naturally, the community was interested in scrutinising the data and to experiment with other motion correction and frame averaging options, and EMPIAR provided the means to publicly distribute the data. EMPIAR-10003 was related to a controversial HIV-1 envelope glycoprotein structure and its deposition enabled others in the field to validate the results (8,22–24). We co-organised the 2016 Cryo-EM Map Challenge, which involved comparing and evaluating contemporaneous processing and validation methodologies using specially selected and curated datasets from EMPIAR including some that were deposited specifically for the challenge (9). EMPIAR continues to play a vital role for the cryo-EM community including for software and methods development and benchmarking, validation/reproducibility, re-analysis, community challenges, and training (25–30). The case for data reuse is well illustrated by EMPIAR-10061 (https://empiar.org/reuse, (31)). As EMPIAR celebrates its tenth anniversary in 2023, it is growing exponentially, **Figures 1 and 2**, having reached 2 PB of archived data (in July 2022) and 1,000 released entries (in September 2022).

**1.**
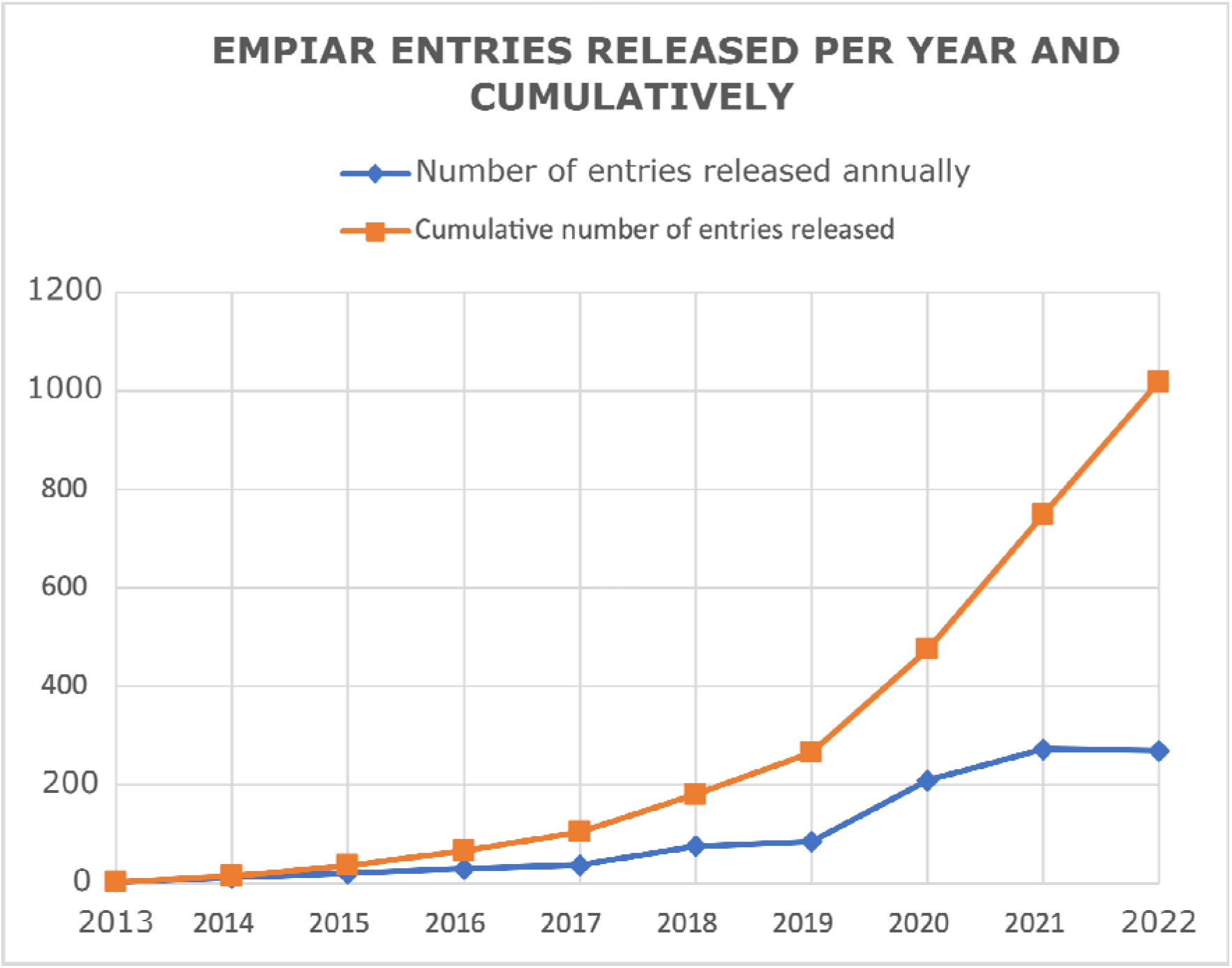
The growth of the number of EMPIAR entries as of the 21^st^ of September 2022 (https://www.ebi.ac.uk/emdb/statistics/empiar_entries_year).

**2.**
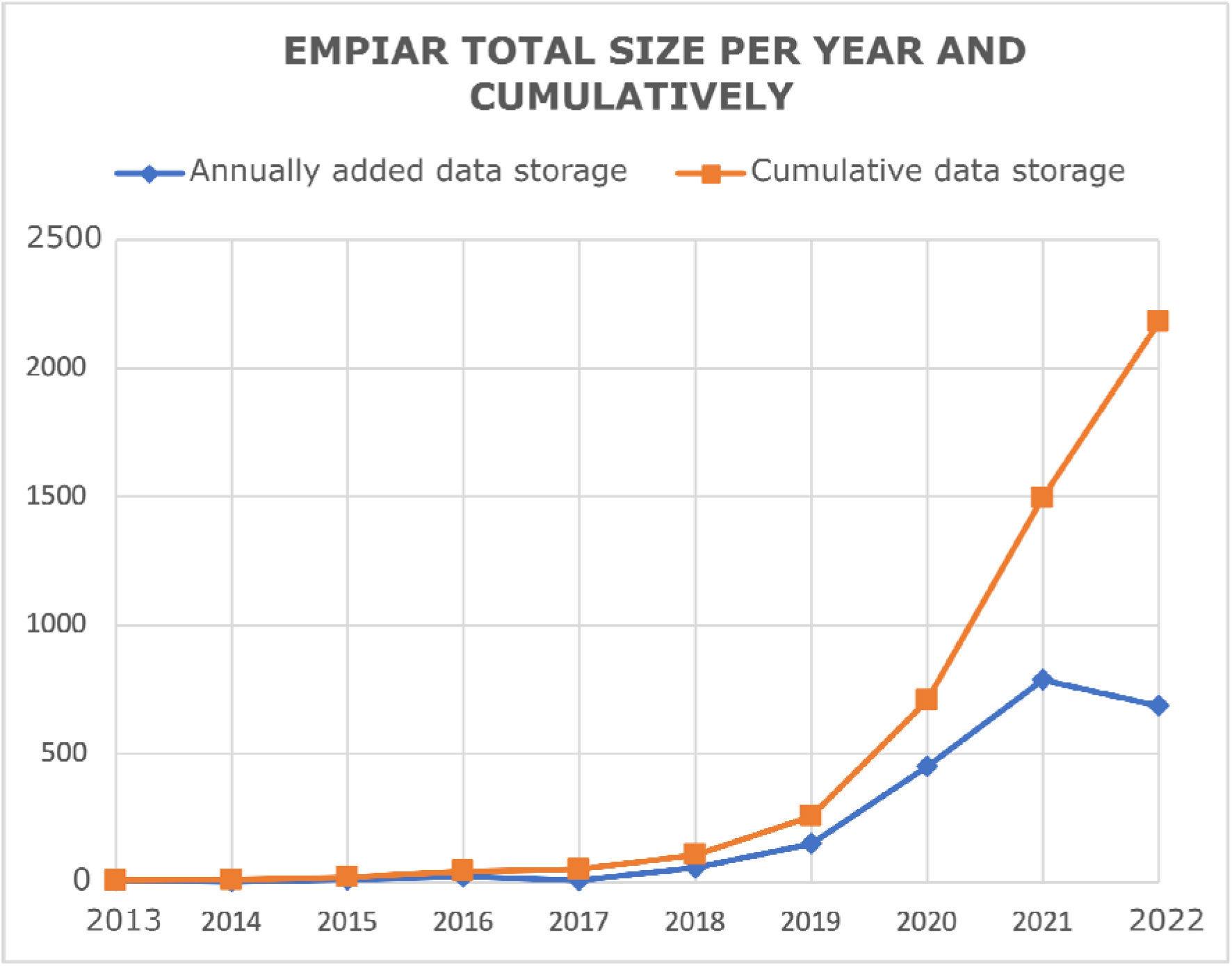
The growth in size of EMPIAR as of the 21^st^ of September 2022 (https://www.ebi.ac.uk/emdb/statistics/empiar_size_increase_year).

In contrast to the cryo-EM community where public archiving of 3D reconstructions in EMDB and fitted or built models in PDB have become well established practises, the notion of public archiving of vEM data is embryonic. It has only gained momentum in the past few years due to an accelerated proliferation of the technique and thanks to leading proponents championing these issues and mustering support within the community. An outcome of the 2012 “A 3D Cellular Context for the Macromolecular World” expert workshop was to expand the remit of EMPIAR to include vEM and X-ray tomography data (17). An expert workshop organised in 2017 considered the issue of public archiving of bioimaging data in general (32) and this led to the establishment of the BioImage Archive (https://www.ebi.ac.uk/bioimage-archive/, (33)) at EMBL-EBI in 2019. The BioImage Archive accepts other forms of imaging data than that archived in EMPIAR, for instance light-microscopy data, thereby also enabling public access to all the data from correlative and multimodal imaging experiments. Together, EMPIAR and the BioImage Archive provide a comprehensive ecosystem for the public archiving of bioimaging data powered by a common IT infrastructure that uses an object store architecture for storage as well as facilities for efficiently transferring large amounts of data to achieve scalability to the petabyte scale and beyond.

## Data deposition (empiar.org/deposition)

Deposition of data to EMPIAR is mainly via a web-based deposition system. Every user of the deposition system has a personal account and different depositors may work together on a deposition with different levels of access assigned to them. For example, the PI and the initial depositor may have full access whereas they may grant read-only access to some coworkers or collaborators to check the entered information. Metadata is entered in a form-based system and includes administrative details such as the names and ORCID identifiers of the depositors, literature references and details of the uploaded image sets. The depositors are asked to upload their data prior to providing the image set metadata. This allows the EMPIAR deposition system to run checks on the data, to analyse the data hierarchy, and to extract metadata from known file types (e.g., MRC-formatted images) which then facilitates and reduces metadata entry for the depositor.

There are two main options that tend to work well to upload large datasets, Aspera (https://www.ibm.com/aspera/connect/) and Globus (https://globus.org). Depositors may also use ftp to upload their data, however due to reliability issues this is an option of last resort, to be used only if the depositor is unable to use Aspera or Globus. With Aspera, the depositor needs to download and install a free web-browser plugin and an application and with Globus the depositor needs to register for a free account, download client software and set up a transfer endpoint. It should be noted that due to the large size of many datasets, data upload may take several days (or even weeks) to complete. While both Aspera and Globus verify the integrity of the upload, a script is provided to the depositor that compares the MD5 checksum of the uploaded data with that of the local copy of the data that was to be uploaded as an optional check. There is also a wrapper script for Aspera and Globus that allows complete or partial deposition based on a pre-generated JSON file with metadata (https://pypi.org/project/empiar-depositor/). The generation of such JSON files has been automated by the EM software package Scipion (http://scipion.i2pc.es/, (34)).

Once all the required metadata has been entered and the data uploaded, the depositor is presented with a button to submit the deposition for further processing. Upon submission the deposition is locked so that the depositor cannot make any further changes and it is passed to the EMPIAR annotation system to be checked by EMPIAR curators. Curators check the deposition using both automated tools, and by manually scanning the entered metadata and data. The annotation system has a communication module (based on Request Tracker (RT, https://bestpractical.com/request-tracker)) that allows depositors and annotators to message one another via email while maintaining a trail in a ticketing system. Many issues that come up during curation can be dealt with by asking the depositor for clarification or further information. In some cases, where data needs to be augmented or replaced, the curator can temporarily unlock a deposition to allow the depositor to make the changes. When the curator is satisfied that the deposition meets EMPIAR data and metadata standards, the depositor will be informed that the data is ready to be released or queued for later release, according to the chosen “release instruction” (see “EMPIAR policies 5.3.1 Release instruction” at empiar.org/policies) and they are asked to confirm that they wish to proceed. If no message is received within two weeks, it is assumed that the deposition has been approved. EMPIAR data is released into the public domain with no rights reserved in accordance with the Creative Commons CC0 designation (https://creativecommons.org/share-your-work/public-domain/cc0/). Depositors can add EMPIAR entries to their ORCiD profiles (https://www.ebi.ac.uk/empiar/orcid).

The release process is initiated by curation staff triggering an automated procedure to move data from a smaller working storage area where active depositions are managed to the EMBL-EBI object store (read-only) where it can be made visible and accessible to the public. It may take several hours or even days from when a release is triggered to when the data is visible to the public due to the large size of some datasets and the time it takes to transfer the data to the object store.

From a user’s perspective, the archive consists of virtual folders, one for each EMPIAR entry. In actual fact there is no hierarchy in the object store but this structure is imposed as a convenience to the user. Each EMPIAR entry folder contains an XML file with the entry metadata in a structure described by the EMPIAR data model (empiar.org/datamodel), and a subdirectory “data” containing all other files. Each image set is a subdirectory in the data directory. For EMDB-related EMPIAR entries most of the metadata is being held by EMDB. For vEM entries EMPIAR will expand its data model to become compliant with recommendations for a basic minimum for metadata collection for imaging data, REMBI (26).

## Data dissemination resources

### EMPIAR website (empiar.org)

The EMPIAR home page provides links, e.g., to the deposition system, a wide range of information including EMPIAR policies and procedures and documentation on accessing the REST API, and several archive statistics and plots. Information boxes provide updates or news affecting the operation of EMPIAR, new resources, recorded tutorials and seminars, etc. A Twitter feed shows the latest tweets from the joint EMDB/EMPIAR account, and the latest EMPIAR-related publications obtained by a keyword search of Europe PMC (35) are also shown.

Central to the page, and its most prominent feature, is a table listing all EMPIAR entries with salient information such as the release date, authors, size of entry and resolution. A simple search system is provided whereby the contents of the table are dynamically filtered as text is typed into the search box. By clicking on entries in the table, the user is taken to the corresponding entry pages and provided with more information and download options. A more sophisticated search system is available as part of EMDB’s combined EMDB/EMPIAR search (https://www.ebi.ac.uk/emdb/search/database:EMPIAR).

### Entry pages (empiar.org/EMPIAR-#####)

Every EMPIAR entry has its own web page which provides summary information, visualisation of images or volumes and the means to download individual images, directories or entire entries. As cryo-EM images typically consist of a movie of large and extremely noisy frames, we precompute thumbnail images by averaging the frames together and down-sampling them so that they are small enough to allow for interactive web-based viewing. The data can be downloaded via Globus (the preferred option), Aspera or as an uncompressed Zip archive streamed via HTTP. The latter option is suitable for smaller datasets typically no more than four gigabytes in size. For all entries with 3D datasets, image data is loaded into the Volume Browser (see below) and links to interactively view these 3D reconstructions are provided at the bottom of the entry page.

### REST API

The REST API (https://www.ebi.ac.uk/empiar/api/documentation) provides programmatic access to EMPIAR metadata in JSON format and is used both internally, for example in the generation of the web pages, and by external resources (e.g., by Scipion (34), https://github.com/scipion-em/scipion-em-empiar). Both GET and POST requests are supported with the latter allowing a range of entries to be specified. Types of calls that are supported include:

- Metadata pertaining to one or more EMPIAR entries.
- Metadata for one or more EMPIAR entries keyed on their corresponding EMDB entry IDs.
- The five latest citations of publications mentioning EMPIAR or EMPIAR entries automatically data mined from Europe PMC.

We provide an API endpoint for journals to check on the status of an entry, e.g. to verify that it is in the system and whether it is released or submitted, e.g., https://www.ebi.ac.uk/empiar/deposition/api/entry_status/EMPIAR-10002/.

### Volume Browser

The Volume Browser (VB; https://www.ebi.ac.uk/empiar/volume-browser) is a web-based resource for interactive visualisation of orthogonal slices through 3D reconstructions from EMPIAR and EMDB. The VB is a further development of and a replacement for the volume slicer shown previously on EMDB web pages as a means of interactively viewing orthogonal slices from a 3D EMDB volume (36). The user may view and move through slices of the 3D reconstruction in any one of the three orthogonal orientations. Controls are provided to quickly switch between the three orientations, to zoom in and out of the reconstruction and to adjust the dynamic range of image voxel intensities. The user can also view a surface reconstruction for related non-tomographic EMDB entries, which can be freely rotated, zoomed and panned. An important feature of the VB is that, unlike its predecessor, it can overlay segmentations onto the 3D reconstruction and present annotations linked to the individual segmentations. The VB landing page features several illustrative examples of volume data overlaid with annotated segmentations. An example is shown in **Figure 3**.

**3.**
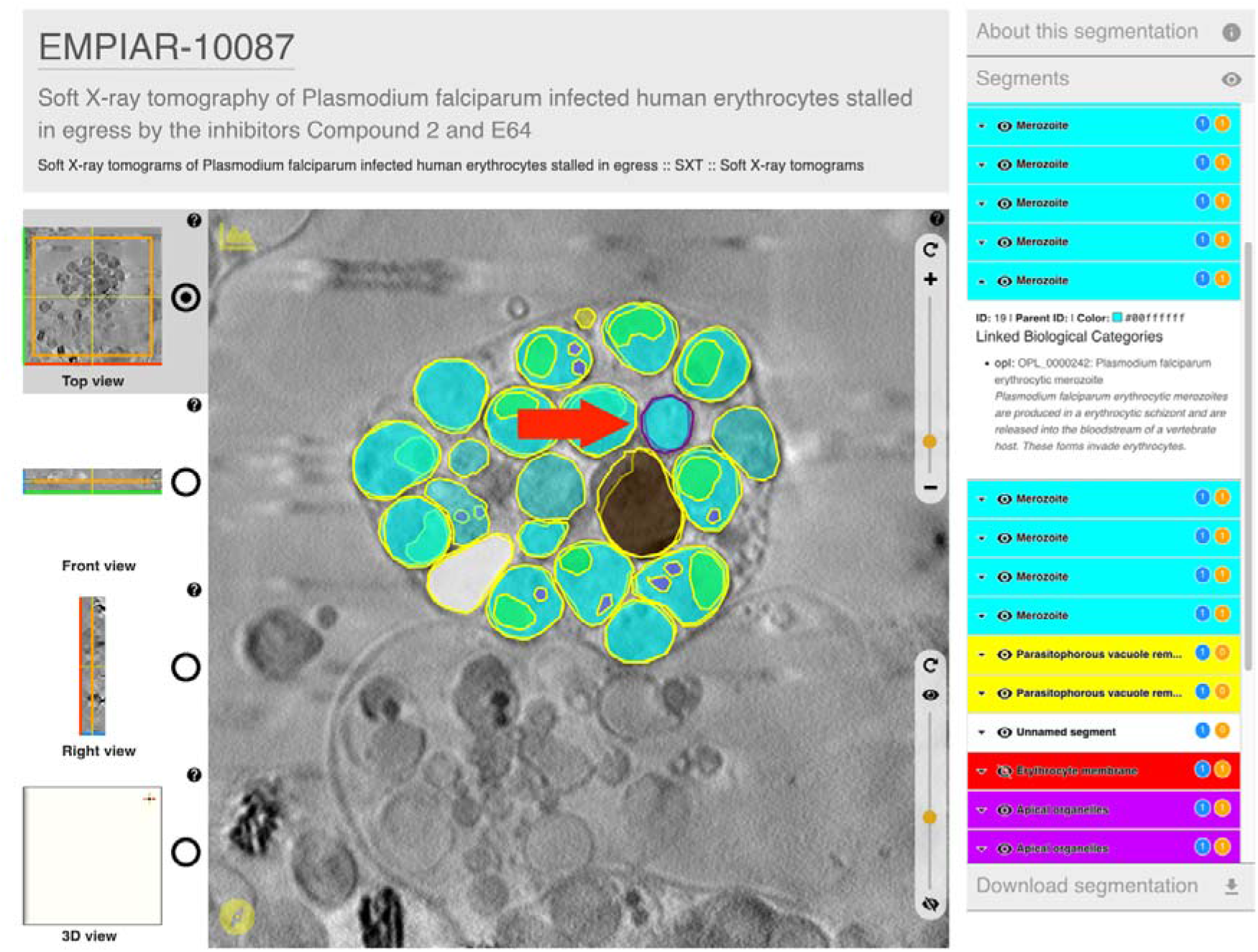
Illustrative example (https://www.ebi.ac.uk/empiar/volume-browser/empiar_10087_e64_tomo03) of an entry in the Volume Browser showing images overlaid with annotated segmentations. Textual annotations are provided in the panel on the right, which displays annotations for the currently selected segment (purple border in the main image panel, indicated by the red arrow).

### Training and social media

An excellent introduction to EMPIAR and the resources available is provided by the EMPIAR Quick Tour (https://www.ebi.ac.uk/training/online/courses/empiar-quick-tour/). For those wishing to delve deeper, especially about depositing data, we recommend the EMPIAR Policies and Processing Procedures document (https://www.ebi.ac.uk/empiar/policies/) and the deposition manual (https://www.ebi.ac.uk/empiar/deposition/manual). The EMPIAR team regularly provide training at courses and workshops and if possible recordings from these are made available on our tutorials page (https://www.ebi.ac.uk/empiar/tutorials/).

We use Twitter (https://twitter.com/EMDB_EMPIAR) to reach out to (potential) users of EMPIAR, **Figure 4**. Weekly tweets focus on particularly interesting entries ranging from biologically important proteins, protein complexes or cells to datasets supporting methods development or obtained with new techniques. Other tweets include announcements of new EMPIAR developments, milestones or job opportunities, and upcoming conferences, courses and workshops. EMDB and EMPIAR share a YouTube channel (https://bit.ly/emchan), which is mostly dedicated to tutorials on how to use the archive, recorded public talks about the resource, featured entries, and highlights of new features.

**4.**
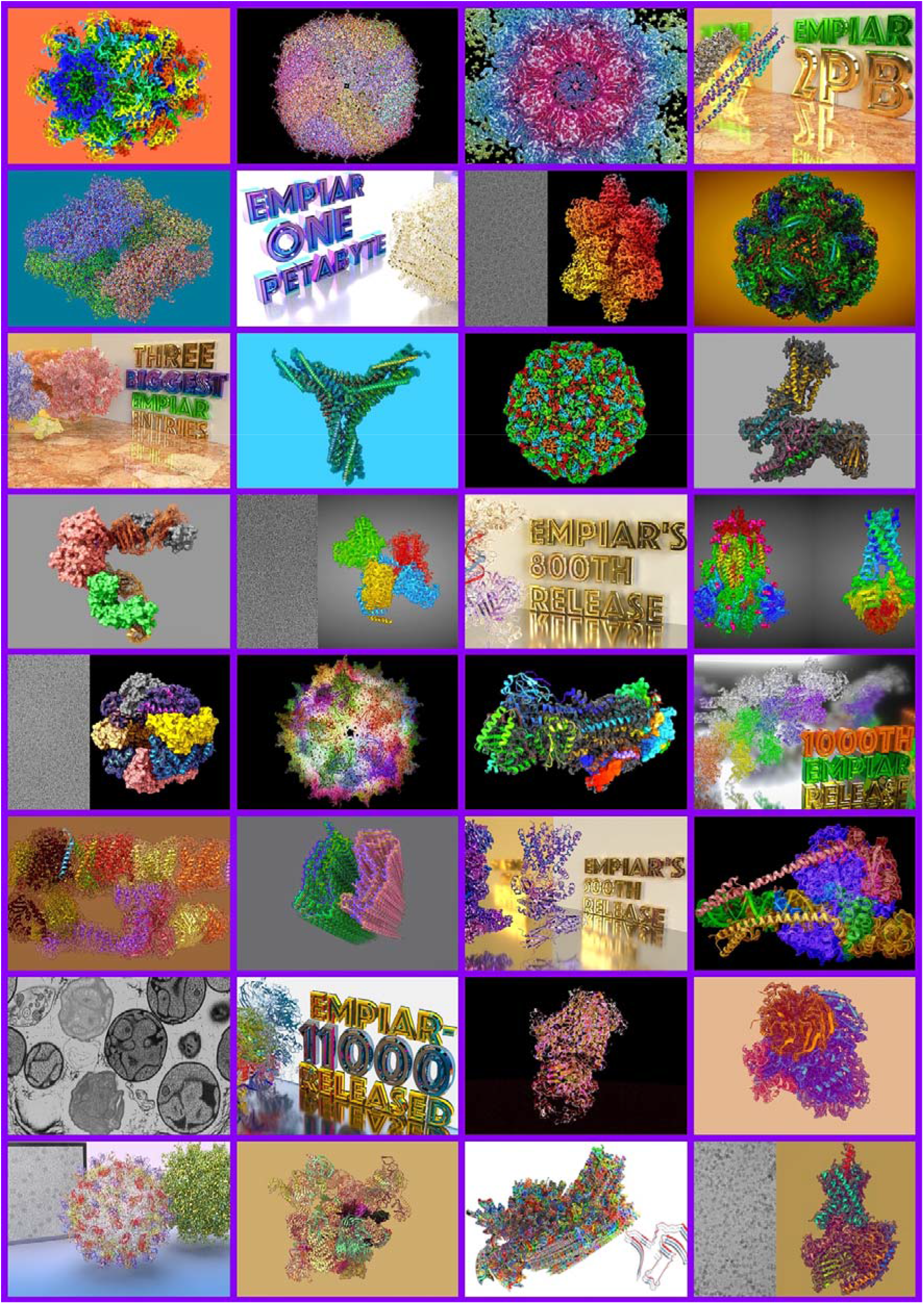
A collage of EMPIAR social media highlights.

## Challenges and outlook

### Expanded metadata collection for volume electron and X-ray microscopy

The metadata currently collected in EMPIAR focuses on describing image sets and not the full experiment. This is satisfactory for cryo-EM experiments where the bulk of the metadata is collected by EMDB but not for vEM and XM experiments where EMPIAR is the sole destination for the experimental data and this severely limits the potential reuse of the data. As a starting point to solving this issue more broadly, REMBI (37) has been defined by the bioimaging community. There is strong support from the vEM community, expressed at the 2021 EMPIAR Volume EM expert workshop, that EMPIAR metadata collection be expanded for vEM to fully support the REMBI specification. Work is currently underway, with the help of the vEM community, to implement this.

### Dealing with correlative imaging data

More and more often, different bioimaging techniques are combined to image a specimen at different scales in order to gain a deeper understanding of what is being imaged. For example, light and electron microscopy can be combined in a correlative light and electron microscopy (CLEM) experiment. The BioImage Archive (33) now provides a home for non-EM and non-X-ray volume imaging data and the EMPIAR deposition system supports linking of entries to any related entries in the BioImage Archive (which in turn links back to the EMPIAR entries). Unfortunately, there are no well-established and accepted standards yet for describing the transformations required to bring different image sets into a common coordinate frame in order to simplify viewing and analysis. Submitters can currently deposit any type of extra correlation-related data or metadata to the BioImage Archive to ensure preservation. Once community standards have been defined, this process will become simpler.

### Data harvesting and pipelining

Manual data entry via web forms is both tedious and error prone. Depositors, archive keepers and end users all stand to benefit from a more streamlined system whereby metadata generated by data acquisition and processing software is read or “harvested” by the deposition system to minimise both manual data entry and the potential errors that go with it. Moreover, the ability to push data from third-party software as the last step in the processing workflow would further facilitate deposition to EMPIAR. We have developed a secure-access API and scripts to enable data harvesting and pipelining. The pipelines have been successfully tested with several facilities, such as EMBL Heidelberg (EMPIAR-10168), the Astbury Biostructure Laboratory in Leeds (EMPIAR-10205) and the National Center for Biotechnology (CSIC) in Madrid (EMPIAR-10514). The latter has integrated the deposition script into their Scipion software and used it to do real depositions. We are also in discussion with developers of various other cryo-EM packages regarding improved harvesting of information generated by their packages, e.g., information about particle coordinates and processing workflows. Properly typed and structured representation of this information in EMPIAR would further enhance the reusability of the data, potentially to the point that the 3D reconstructions deposited to EMDB could be automatically reproduced from the image sets deposited to EMPIAR.

### Archiving of segmentation data

At the resolutions typically achieved in vEM and cryo-ET experiments, segmentations and their biological annotations play a vital role in enabling the (re)use of the data by biologists and bioinformaticians and it is therefore important that segmentations are also publicly available. Currently, segmentations may be deposited to EMPIAR and EMDB but only as part of a data submission by the original data depositors. However, there are many instances where segmentations have not been performed by the same team that generated the data. In some cases, the enormity of the task has led to outsourcing of the segmentation to specialist groups analysing different targets or regions or employing different methodologies. A recommendation of a 2021 EMPIAR vEM workshop was to allow segmentations to be deposited to EMPIAR as entries in their own right with links maintained to related EMPIAR data sets. This would accommodate both scenarios of segmentation provenance.

### Dealing with data growth

Unlike the deposition of atomic models to PDB and 3D reconstructions to EMDB, which are now mandatory for publication in most journals publishing structural biology studies, the deposition of data to EMPIAR is generally not mandatory for either cryo-EM or vEM publications. Typical cryo-EM datasets are in the terabyte range (currently around 3 TB on average); as of September 2022 there are 38 datasets (~3.7% of the total datasets) exceeding 10 terabytes in size. Typical vEM depositions to EMPIAR are much smaller, usually in the 10-100 GB range, but there are a few large projects and institutes, e.g., in the area of connectomics, that generate data on the petabyte scale. An important question that needs to be addressed is therefore how EMPIAR infrastructure would cope if deposition of data for cryo-EM and vEM were to be made mandatory.

In terms of storage, EMPIAR relies on the EMBL-EBI object store which in technical terms is well scalable to the exabyte range. The cryo-EM community has benefitted greatly from this hot storage, making it possible to download tens of terabytes of data at the click of a button. However, the storage and IT infrastructure is funded by the UK government and the UK Research and Innovation Strategic Priorities Fund (UKRI SPF) data infrastructure program and sustained future investment relies on our ability to justify that this is money well spent. We are therefore considering adding metadata to EMPIAR entries to track lifecycle events, e.g., date of last access, number of downloads etc. With this information it will be possible to better manage storage of the data, for instance by allowing data that is older and rarely accessed to be moved onto “colder” but cheaper storage. The large-scale vEM projects mentioned above tend to have significant resources to host and provide access to their data long term. In a 2021 EMPIAR Volume-EM workshop it was proposed that for such large datasets, rather than duplicating the data in the centralised store, EMPIAR could maintain links to the project site and only store relevant metadata and possibly a final condensed version of the 3D reconstruction that could be used for viewing purposes.

In addition to increased storage, an increase in the number of depositions will also pose challenges in terms of the curation of the data. We have a collaboration with PDBj (Osaka, Japan) who maintain a mirror of EMPIAR (https://empiar.pdbj.org/) and who are also data brokers for Japanese depositions, i.e., they assist Japanese depositors in preparing depositions which are then pushed to EMPIAR. As EMPIAR continues to grow, it may be necessary to consider further collaborations with regional partners in order to have a sustainable solution for the public archiving of EM data.

### A FAIR analysis of EMPIAR data

Wilkinson *et al*. outlined the FAIR (Findable, Accessible, Interoperable and Reusable) guiding principles for good data management and stewardship by public archives (38) and it is useful to gauge where EMPIAR stands with respect to these aims. Every EMPIAR entry has a persistent identifier and meta-data indexed in a searchable resource, which is needed to make the data findable. As discussed earlier, work is underway to substantially enhance the richness of the metadata and this should further improve the findability of the data. All EMPIAR data is freely and openly accessible via a range of download options provided at no cost to the user. However, the very size of these datasets may hinder accessibility and technological solutions (such as data compression or the ability to download parts of datasets at the desired sampling) may be required to make the data accessible to a wider range of users. The formal specification of the EMPIAR metadata model as an XML schema aids interoperability. This is further enhanced by the initiatives in place for sharing and linking information with EMDB and the BioImage Archive, the incorporation of relevant references to these archives and the eventual use of REMBI to underpin the vEM data model. Despite the numerous examples of reuse of EMPIAR data, we know anecdotally that the lack of sufficient metadata describing the data is impeding wider reuse of EMPIAR, e.g., for automated reprocessing of datasets and as training data for machine learning approaches. In summary, improving the richness of the metadata description of the data is fundamentally important to maintain and increase the relevance and usefulness of the archive and is therefore a key focus of future development.

## Acknowledgements

We thank José Salavert-Torres and Carlos Lugo Vélez for their contributions to the early stages of development of EMPIAR. We thank Matthew Hartley (BioImage Archive Team Leader; EMBL-EBI), Mary Barlow (Head of Campus Ops & Capital Projects; EMBL-EBI), Andy Cafferkey (Technical Services Team Leader; EMBL-EBI) and their respective teams for their help and collaboration in building up and maintaining EMPIAR, the BioImage Archive and the underpinning IT infrastructure. We thank Prof. Genji Kurisu and the PDBj team for the collaboration in developing EMPIAR-PDBj. Finally, we would like to thank the very many members of the EM community who have enthusiastically deposited data, participated in workshops and provided help, advice, and feedback.

